# At-home, cell-free synthetic biology education modules for transcriptional regulation and environmental water quality monitoring

**DOI:** 10.1101/2023.01.09.523248

**Authors:** Kirsten J. Jung, Blake J. Rasor, Grant A. Rybnicky, Adam D. Silverman, Janet Standeven, Robert Kuhn, Teresa Granito, Holly M. Ekas, Brenda M. Wang, Ashty S. Karim, Julius B. Lucks, Michael C. Jewett

## Abstract

As the field of synthetic biology expands, the need to grow and train science, technology, engineering, and math (STEM) practitioners is essential. However, the lack of access to hands-on demonstrations has led to inequalities of opportunity and practice. In addition, there is a gap in providing content that enables students to make their own bioengineered systems. To address these challenges, we develop four shelf-stable cell-free biosensing educational modules that work by just-adding-water and DNA to freeze-dried crude extracts of *Escherichia coli*. We introduce activities and supporting curricula to teach the structure and function of the *lac* operon, dose-responsive behavior, considerations for biosensor outputs, and a ‘build-your-own’ activity for monitoring environmental contaminants in water. We piloted these modules with K-12 teachers and 130 high school students in their classrooms – and at home – without professional laboratory equipment or researcher oversight. This work promises to catalyze access to interactive synthetic biology education opportunities.

## Introduction

Incorporating hands-on, active learning teaching methods at the secondary and undergraduate school levels increases student retention and performance across science, technology, engineering, and mathematics (STEM) disciplines [1-3]. In particular, biotechnology lab-based research, often done in high schools through inter-institutional joint studies such as the *Wolbachia* Project or SEA-PHAGES program [4, 5] contributes to significant improvement in students’ self-reported technical and professional skills [6, 7].

Educational laboratory modules for synthetic biology, the fundamental science and engineering research that engineers biology to tackle global challenges (e.g., access to medicines, sustainable manufacturing), [8, 9] are in particular demand. Indeed, the acceleration of private and public investment in synthetic biology, [10-13] as well as the recent United States Presidential Executive Order on Advancing Biotechnology and Biomanufacturing [14] has heightened the need for supporting student literacy and learning. Such education is essential for long-term field participation, civic engagement, and ethical development [15].

Rapid growth of organizations like the International Genetically Engineered Machines (iGEM) [16, 17] competition, and the availability of do-it-yourself experiments from BioBuilder [18], Amino Labs (https://amino.bio), and the ODIN (https://www.the-odin.com) highlight this potential [19]. However, the high cost, slow pace, and regulatory bottlenecks and biocontainment requirements for experiments using engineered microbes preclude hands-on participation of many aspiring synthetic biologists [9], particularly students in under-resourced communities and schools.

Recently, cell-free gene expression (CFE) [20, 21] has emerged as a useful platform for widening access to synthetic biology education. In CFE systems, protein synthesis is carried out by transcription and translation machinery (polymerases, ribosomes, tRNAs, etc.) that has been harvested from living cells and reconstituted in a test tube [22]. The preparation of extract from *Escherichia coli* cells has been extensively optimized to allow for: high protein synthesis yields [23, 24]; on-demand synthesis of conjugate vaccines [25, 26], antibodies and antibody-drug conjugates [27-29], antibody fragments [30, 31], and antimicrobial peptides [32, 33]; detection of toxic metals and organic pesticides in drinking water [34-38] or nucleic acids in biological samples [39-43]; and rapid prototyping of genetic parts [44], enzyme pathways [45, 46], and post-translational modifications [47-49] using high-throughput liquid-handling robotics.

Cell-free expression systems are particularly suitable for decentralized distribution in low-resource settings, including classrooms, because the reactions are stable in a lyophilized format for months at room temperature [50]. Consequently, protein synthesis can be initiated just by adding water and a template DNA [41]. Using freeze-dried CFE reactions, we and others have developed experimental learning modules for high school and undergraduate students, focused on teaching the central dogma [51-53], antibiotic resistance [54], and the mechanism of CRISPR-Cas9 [54, 55]. The simplest of these modules has been commercialized as the BioBits^®^ Central Dogma kit [56].

Early successes with cell-free education kits nevertheless left several gaps to be addressed. First, because students performed most of the published experiments in well-equipped university labs rather than a high school classroom, widespread access was not guaranteed [51, 54, 55]. The importance of minimal-equipment experiments was made even more apparent by the COVID-19 pandemic, during which many students were unable to attend a classroom. Second, previous work demonstrating student success in the experimental modules was limited to small sets of students and failure modes were not outlined, either for experimental scaleup or for individual student performance. Finally, the proposed experiments did not allow students the creative freedom to design, build, and test their own engineered systems. In a world where the half-life of cutting-edge technology is decreasing, students need learner-led opportunities that afford them the ability to evaluate new material and test potential answers (or at least organize the possibilities) to unsolved problems.

Here, we set out to develop freeze-dried, cell-free educational modules that could address these gaps to facilitate both inquiry-based learning and at home usage for expanded access and impact. As a model, we focused on biosensing. Three experimental modules allowed students to interrogate the mechanism of the regulation of the lactose-inducible operon from *E. coli*, measure the dose-response of the LacI repressor protein to isopropyl β-d-1- thiogalactopyranoside (IPTG; lactose analog), and compare the performance of four reporter outputs. To facilitate inquiry-guided learning, we also developed a fourth “design-your-own-biosensor” module, leveraging cascaded genetic circuitry to design, build, and test cell-free biosensors for the detection of toxic contaminants in drinking water. Finally, we investigated the distribution of cell-free education modules at scale. We prepared kits for remote use by 130 advanced high school biology students (including several who performed the experiments in their own homes). Overall, the student experimental success rate varied between 60 and 100%, depending on the module and scale. We measured significant improvement in student understanding and engagement after completing the modules, and also identified stumbling-blocks for further scaling out this technology to more schools, teachers, and scientists-in-training.

## Results and Discussion

### Preliminary design of a biosensing education kit

We set out to create cell-free biosensing education modules that could enable students to build their own cell-free biosensors and be carried out in a distributed, at home setting. To start, we designed three simple experiments to explore the design of a cell-free biosensor. We chose the *E. coli* lac operon as a model since it is widely taught as an example of a negative-inducible system [57]. The allosteric transcriptional factor LacI represses transcription initiation at a lactose-inducible promoter (pLac) containing the operator sequence (*lacO*) (**Figure 1A**) [58, 59]. We built a lactose-inducible reporter plasmid in which the coding sequence for monomeric red fluorescent protein (mRFP) is placed downstream of the *lacUV5* promoter. Then, as previously described [34], we pre-enriched an *E. coli* cell extract with LacI by overexpressing the transcription factor during growth (**Figure 1A**).

**Figure 1.**
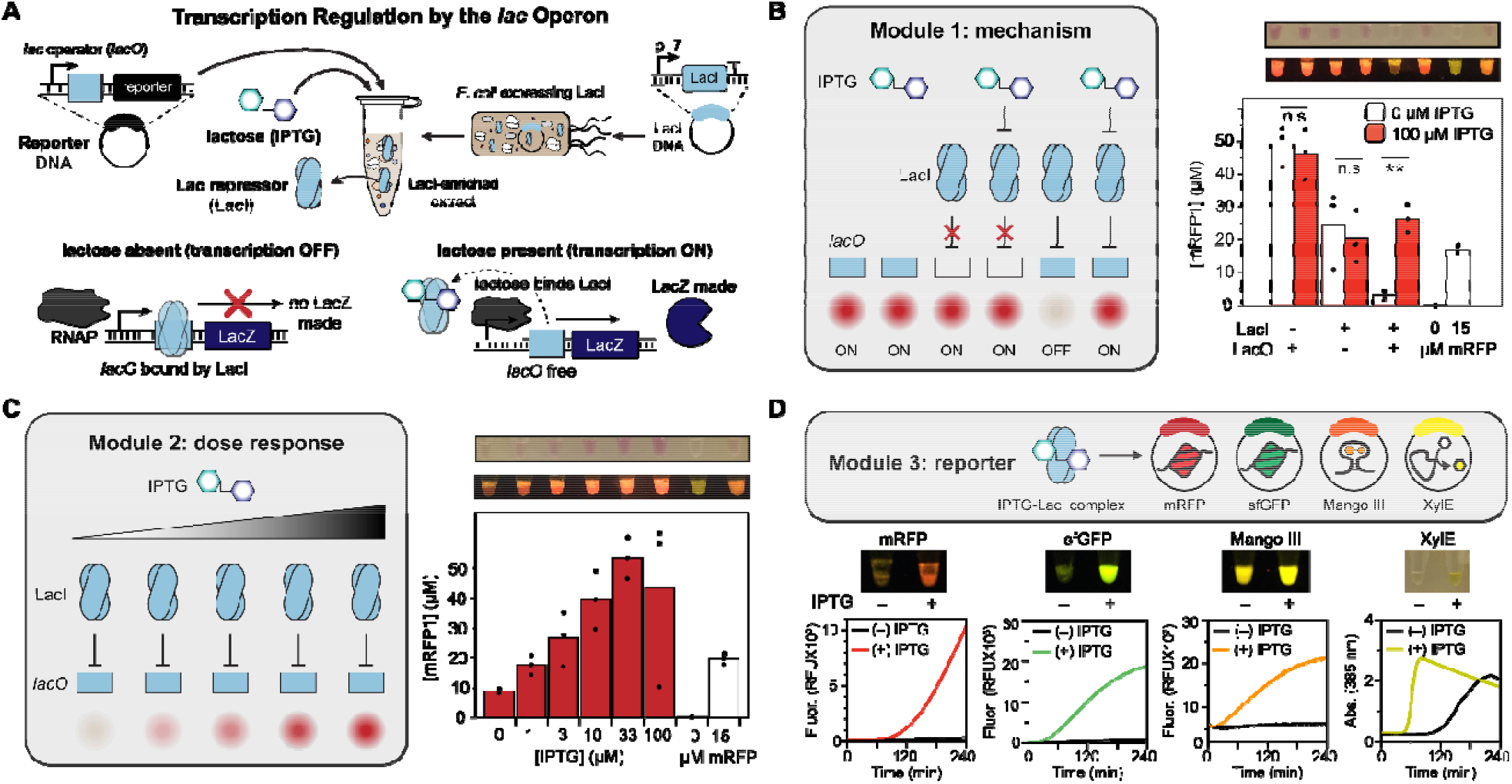
Design of a synthetic biology education module for transcription regulation by the lac operon. (**A**) Overview of cell-free lactose sensor. The Lac repressor (LacI) is over-expressed in *E. coli* used to prepare source extract and represses its target promoter, containing a copy of the lac operator sequence upstream of the reporter protein or RNA gene. (**B**) Laboratory data for a module designed to teach the mechanism for the Lac repressor using an eight-strip of PCR tubes. Tubes 1 and 2 contain lyophilized CFE reactions without LacI (*i*.*e*., only blank extract) and 40 nM of the reporter plasmid pLac-mRFP1. Tubes 3 and 4 contain lyophilized reactions with LacI, but the reporter plasmid lacks a *lac* operator (J23119-mRFP1). Tubes 5 and 6 contain the full sensor (LacI-enriched extract and pLac-mRFP). Tubes 7 and 8 are controls containing, respectively, 0 and 15 µM of purified mRFP1. Upon rehydration with 20 µL either water (tubes 1, 3, 5, 7, and 8) or 100 µM IPTG (tubes 2, 4, and 6), repression is only observed in tube 5, where the repressor and operator are present and no ITPG is supplied. Plotted data represent the average and individual endpoint yields of mRFP1, computed from a fluorescence calibration curve generated by purified mRFP1, from three independent technical replicates for the full eight-strip module, rehydrated in a research laboratory and incubated at 30 °C for 24 hours. Sample images of one replicate are shown in white and blue light (using the miniPCR bio™ P51(tm) Molecular Fluorescence Viewer) from an iPhone photograph. ** represents p < 0.05; n.s. indicates no significant difference between the + and – IPTG conditions. (**C**) Laboratory data for a module designed to teach the dose response behavior of the LacI sensor. Freeze-dried reactions containing LacI and 40 nM pLac-mRFP1 were rehydrated with the indicated concentration of IPTG, and incubated at 30 °C overnight, then quantified by plate reader. The 0 and 15 µM mRFP controls were maintained in this module. (**D**) Laboratory data for a module designed to teach the advantages and disadvantages of alternative reporter outputs for the LacI sensor. Freeze-dried reactions were prepared with LacI-enriched extract, half containing 0 µM IPTG (tubes 1, 3, 5, and 7), and half containing 100 µM IPTG (tubes 2, 4, 6, and 8). The lyophilized reactions were rehydrated with 20 µL of 40 nM pLac-mRFP1 plasmid (tubes 1 and 2), 20 nM pLac-sfGFP plasmid (tubes 3 and 4), 80 nM pLac-Mango III plasmid + 2 µM T01:biotin (RNA aptamer, tubes 5 and 6), or 10 nM pLac-XylE plasmid + 2 mM catechol and pipetted onto a plate reader to measure the kinetics of sensor activation over four hours at 30 °C. Colored lines represent the average of three (+) IPTG conditions; black lines indicate (-) IPTG. Photos indicate reaction yield at the endpoint in blue light (fluorescent outputs) or white light (XylE).

For the Module 1 experiment (mechanism), we prepared three reaction conditions, each comprising 20 µL cell-free sensors with all requisite CFE reagents, freeze-dried with the extract and reporter plasmid. The first pair of 20 µL reactions included 40 nM of the pLac-mRFP plasmid, but no pre-enriched LacI beyond the genomic copy in our BL21 Star (DE3) *E. coli* chassis strain. In the second pair of reactions, 5% of the reaction’s unenriched extract was replaced with an extract pre-enriched with LacI, a fraction that we found was optimal for repression (**Supplemental Figure S1**). However, transcription from the provided reporter plasmid (J23119-mRFP) is constitutive because this promoter lacks an operator site. The third pair of reactions constituted the intact sensor, with both LacI present in the extract and a pLac-mRFP reporter plasmid. We also included positive and negative mRFP calibration controls consisting of lyophilized purified protein at a concentration of 0 or 15 µM. Thus, the full module constituted an eight-strip of PCR strip tubes (**Figure 1B**). When rehydrated with either 0 or 100 µM IPTG inducer, only the third pair of tubes showed both repression and induction: without either the repressor or operator, the sensor was broken and constitutively ON. Conveniently, at a 24-hour endpoint, sufficient mRFP was synthesized in the ON conditions to be visible to the naked eye or under a blue light imager [51].

Following these experimental controls, Module 2 (dose response) was then designed with the aim of teaching students about the concentration dependence of biosensors (simplified, to omit the effects of catabolite repression and cooperativity [60]). Six tubes containing the inducible lactose sensor (containing 5% LacI extract and the pLac-mRFP plasmid) were freeze-dried along with the same calibration controls, and the reactions were rehydrated with increasing concentration of IPTG. As expected, a smooth increase in red fluorescence was observed, up to saturation at 100 µM (**Figure 1C**).

Finally, in Module 3 (reporter), we explored the choice of output for a putative point-of-use biosensor. In addition to mRFP, we cloned superfolder green fluorescent protein (sfGFP) [61], the RNA aptamer Mango III [62, 63], and the colorimetric enzyme XylE (catechol 2,3-dioxygenase [35]), under transcriptional regulation by the pLac promoter. After optimizing the concentration for each reporter DNA (**Supplemental Figure S2**), we induced activation of each reporter in the presence of IPTG, though with different degrees of sensor leak/background signal and time to result. Predictably, the RNA aptamer was the quickest reporter to be produced, but it was invisible to the naked eye, and its background fluorescence from the T01:biotin dye was high under blue light. Also as expected, sfGFP folded faster than mRFP and generated a signal at an earlier time point; both fluorescent proteins could be observed at the endpoint under white and blue light. XylE activated quickly (measured by the development of yellow color under white light), but it also had the greatest amount of leak due to the enzymatic turnover and unavoidable transcriptional leak through LacI, as previously described (**Figure 1D**) [64].

### Small-scale implementation of three biosensing education modules in high school classrooms

Next, we adapted Modules 1-3 for a classroom setting. We developed a roughly one-week-long curriculum that guided students through the three experiments and developed curricula intended for an advanced secondary school (AP Biology) classroom (all curricula included in **Supplemental File 6**). We then prepared 25 freeze-dried reaction strips for each module, identical to the experiments performed by trained researchers in **Figure 1**. These experimental modules were then performed by 22 AP Biology students in their own classrooms at Evanston Township High School over the course of a week. To simplify the experiment and remove some sources of external error, all students had been previously exposed to micropipettes and cell-free gene expression through the BioBits^®^ Central Dogma kit.

For Modules 1 and 2, the rehydrated reactions were incubated at room temperature for 48 hours in the classroom and then transported back to our laboratory for image analysis and quantification by plate reader. For Module 3, which was time sensitive, we instead captured images of the rehydrated tubes in white and blue light initially, after 1 hour, and after 24 hours, on-site in the classroom. The resulting experimental data and representative photographs are plotted in **Figure 2**. The full data set from students, including uncropped photographs, are available in **Supplemental Files 1-3**.

**Figure 2.**
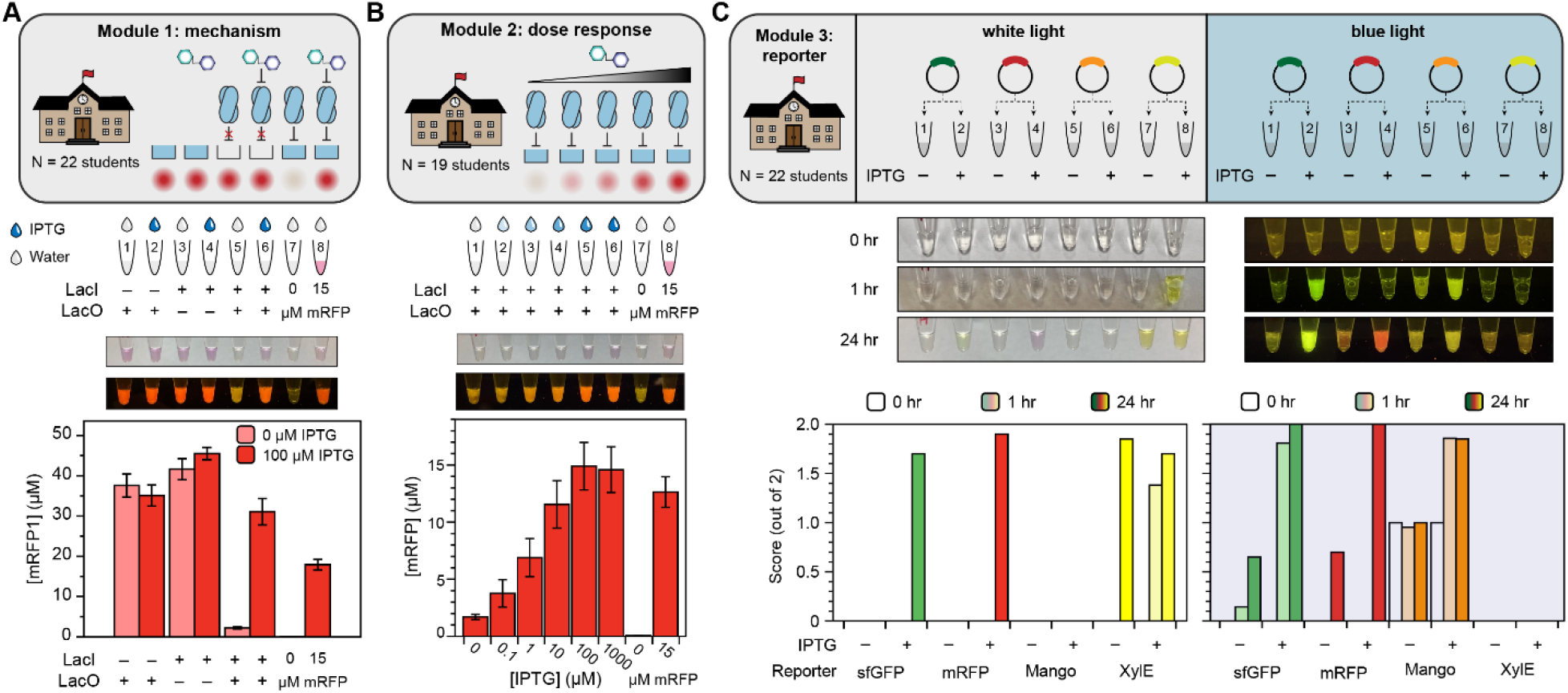
Results of small-scale deployment of biosensing educational kit in advanced high school classroom. (**A**) Implementation of Module 1 (mechanism) experiment in high school classroom. Students added either water or 100 µM IPTG to the indicated tubes using a micropipette, and the reactions were incubated for 48 hours at room temperature. Then, the reactions were collected, and yields were quantified by plate reader. Bars represent the average, and error bars represent the standard error of the mean, from 22 replicates. Inset photos represent white and blue light photos (in P1 imager) of a sample replicate. (Full student data for all modules are available in the **Supplemental Files**) (**B**) Implementation of Module 2 (dose response) experiment in high school classroom. Reactions were incubated for 48 hours at room temperature and quantified by plate reader. Data represent the average and standard error of the mean from 19 student replicates, with one representative sample photo taken at endpoint in white and blue light. (**C**) Implementation of Module 3 (reporters) experiment in high school classroom. Students added reporter plasmids as indicated; then, reactions were incubated at 30 °C and photographs were taken in white and blue light at t = 0 hours, 1 hour, and 24 hours for each reaction set. Researchers then qualitatively assigned to each photograph values 0, 1, or 2 to represent “OFF”, “FAINT”, or “ON”, respectively. The indicated bars represent the average score from 21 student replicates of the reporter activity at each time point (indicated by bar shading), for each reporter (indicated by bar color), and in white and blue light (left and right plots, respectively). One sample time-course with paired photos is shown.

The students’ data generated in the classroom matched the laboratory data very well. Dispensing either water or IPTG solution from micropipettes, the students generally achieved the expected qualitative results in Modules 1 and 3; somewhat greater variability was observed Module 2, possibly due to errors in serial dilution (the students performed their own dilutions from a 100 µM IPTG stock). Of the 22 students who performed Module 1, 16 had a “perfect” response of ON, ON, ON, ON, OFF, ON, OFF, ON for tubes 1-8, and over 90% of the individual tubes matched the expected behavior. The experiment only failed for one student. Thirteen of the nineteen students who performed Module 2 also observed the expected smooth, monotonic increase in mRFP production between tubes 1-6 as IPTG dose increased.

To quantify success rates in Module 3, since we could not take plate reader measurements at intermediate timepoints in the classroom, we instead assigned qualitative brightness scores of 0, 1, or 2 to each reporter, with and without the inducer, at each timepoint, based on the students’ photographs in white and blue light. After one hour, most students observed the sfGFP and Mango reporters to be ON in blue light, and the XylE reporter was ON in white light. Increases in both signal and leak were observed for all reporters after 24 hours. In many cases, the XylE signal was stronger in the OFF state than the ON state after 24 hours because the product of the chemical reaction degrades.

### Evaluation of educational impact of biosensing kits at secondary school level

To assess whether the biosensing kits were effective in inspiring and teaching the students who did the activities in the small-scale implementation, we measured how well the program goals were met through pre- and post-module surveys (**Figure 3A**). The survey consisted of a series of statements and asked the student to rate whether they agree with the statement or not on a scale from 1-5, with a score of 1 indicating that they strongly disagree with the statement, a score of 3 indicating that they neither agree nor disagree, and a score of 5 indicating that they strongly agree with the statement. Prior to completing any of the three experimental activities, students were asked to fill out a survey to establish baseline biology knowledge and perceptions. Following each of the experimental activities, but before seeing any material for the next activity, students were asked to take the same survey again to capture changes as a result of participating in that module.

**Figure 3.**
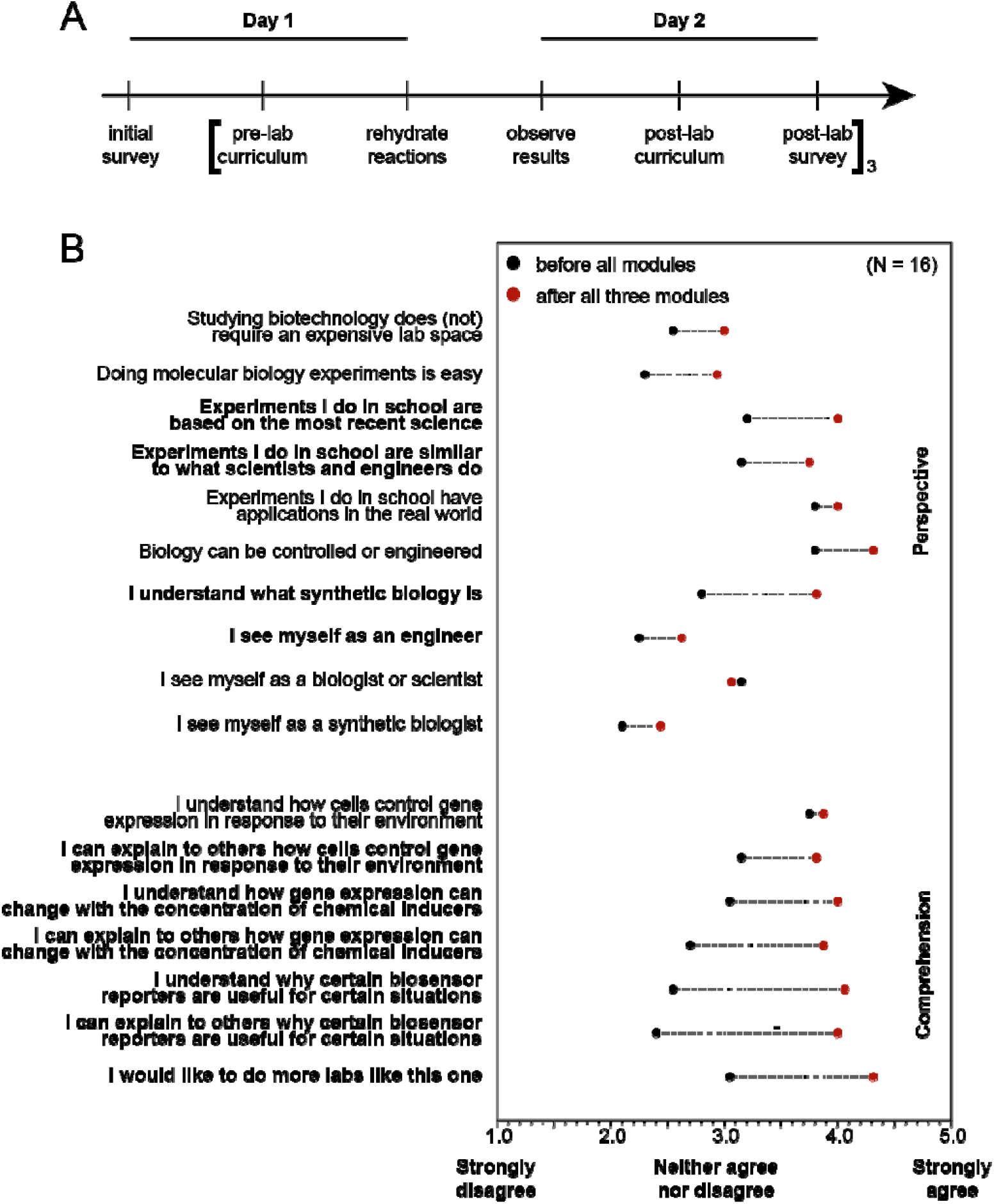
Experiential impact on students’ learning from biosensing modules. (**A**) Sample timeline for performing Modules 1 through 3 in a week of experimental instructional learning, including pre- and post-lab surveys. (**B**) Survey results from 16 students who performed all three experimental modules. Black dots indicate the average scores (between 1 and 5) to the “comprehension” and “perspective” questions before performing the experiments, and red dots indicate scores afterwards. Bolded questions showed a significant (p < 0.05) increase in score across paired student replicates from a Wilcoxon signed-rank paired test. Questions match exactly what was present in the student surveys, with the exception of the first statement which was written in the negative form, “Studying biotechnology requires an expensive lab space”, and has been negated here for clarity.

Overall, survey questions were designed to assess two outcomes of the experimental activities: (1) changes in perspective of science and engineering and (2) changes in comprehension of biological and engineering concepts. Generally, participation in the three-activity series increased survey scores across both categories (**Figure 3B**). Participation in the activities significantly increased how likely students were to agree with the perspective statements that: the experiments that they do in school “are based on the most recent science” and are “similar to what scientists and engineers do”, as well as that they “understand what synthetic biology is”, that they “see themselves as engineers”, and that they “would like to do more experiments like this one.” Likewise, participation in the activities significantly increased how likely students were to agree with the comprehension statements that they understood and could explain the material.

### Development of an advanced module for biosensor design

We developed the initial three experimental modules to teach regulation by the *lac* operon because it is a well-understood biosensor in *E. coli*. However, towards the goal of engaging students in synthetic biology, and the known learning benefits of having students solve problems, answer questions, and formulate questions of their own [1, 65, 66], we next created an avenue for students to design, build, and test their own cell-free biosensors. As a model, we developed a biosensor activity to detect water contaminants of public health concern.

Previous efforts to engage students in the synthetic biology design-build-test-learn framework have struggled with the build phase due to challenges in DNA assembly and transformation(roadblocks that hold back many iGEM teams as well [67]). Cell-free expression circumvents some of these challenges: PCR-amplified linear DNA can be used in lieu of cloned plasmids, and transformation is unnecessary. However, as a different approach, we decided to use a cascaded amplifier circuit [68] to decouple the genetic linkage between the sensing elements (allosteric transcription factor (aTF) and inducible promoter) and the reporter protein. Briefly, the cascaded amplifier uses an orthogonal T7 RNA polymerase (o-T7 RNAP) as the output of the inducible promoter, and a second plasmid encodes the reporter protein under the control of the corresponding T7 promoter (P_o-T7_) (**Figure 4A**). The advantage of this setup is that we could lyophilize all sensor elements together (the aTF-enriched extract and its corresponding inducible sensor plasmid, pReg-o-T7 RNAP) to make an arbitrary ligand-sensing reaction, but without a defined transcriptional reporter. Then, to simultaneously build and test the sensor, students could rehydrate the reactions with liquid stocks containing the desired reporter plasmid and any co-substrates: essentially, running any or all of the Module 1-3 experiments as desired for environmentally relevant contaminants, with a single set of reporter plasmids.

**Figure 4.**
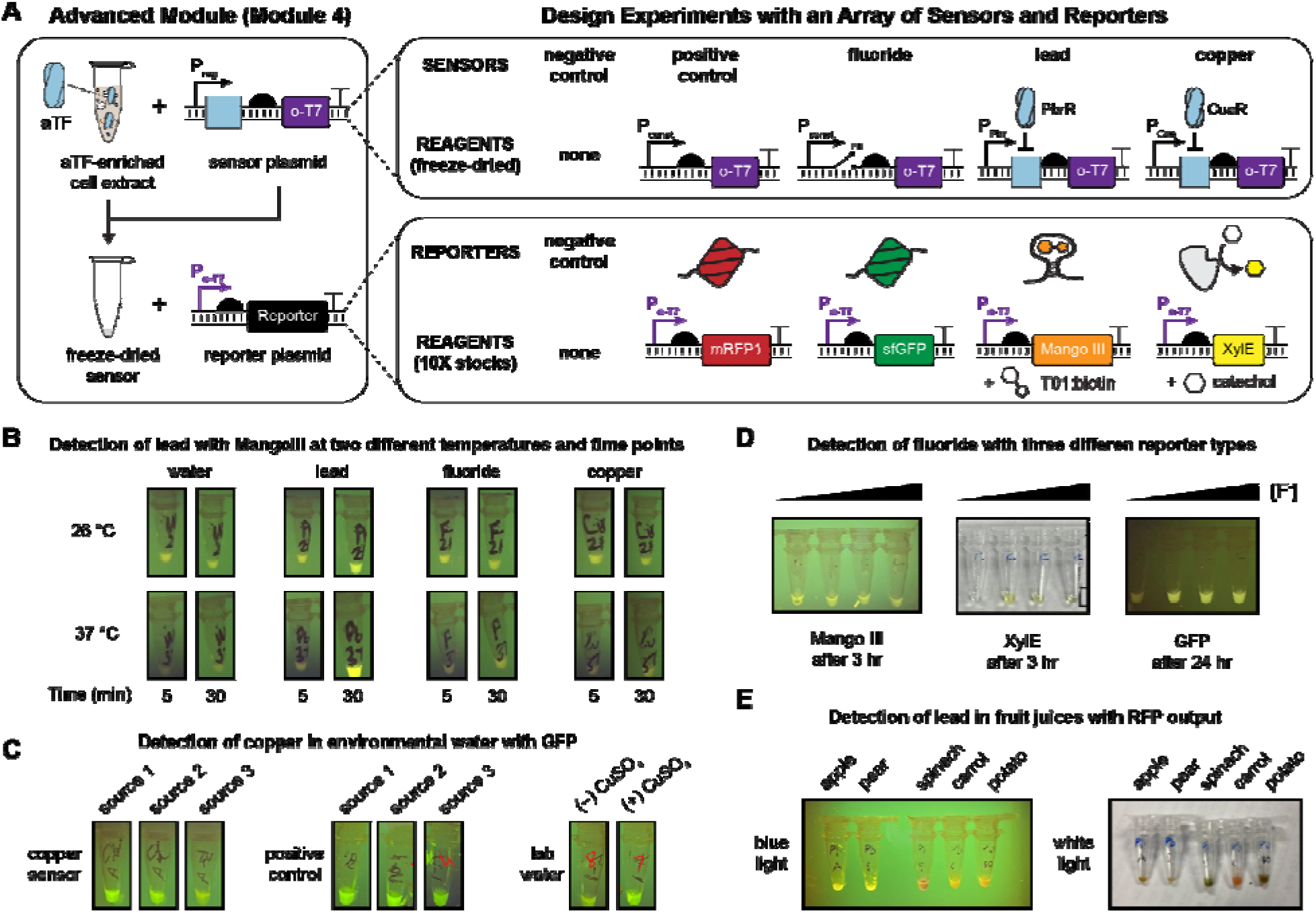
Advanced experimental module for student-initiated biosensor design. (**A**) Overview of Module 4 (advanced design). Students were provided freeze-dried sensor reactions containing cell-free extracts pre-enriched with the allosteric transcription factors PbrR (lead sensor) or CueR (copper sensor) as well as the respective sensor plasmids, in which a regulated, paired promoter (pPbr, pCue), riboswitch (fluoride sensor), or constitutive promoter (positive control) drives expression of orthogonal T7 RNAP in an RNA polymerase cascade. (A negative control sensor reaction lacked a sensor plasmid to synthesize o-T7 RNAP.) They were also provided, in 10X concentrates, plasmids encoding the Module 3 reporters and substrates mRFP1, sfGFP, Mango III + T01:biotin, and XylE + catechol, with the reporters placed under control of the o-T7 promoter. Finally, they were given liquid stocks for copper, lead, and fluoride. By rehydrating the sensor reaction with a reporter plasmid and analyte of choice, students could therefore build, test, and design arbitrary sensor-output pairs for common inorganic water contaminants—all without any cloning. (**B-E**) Sample results from four experiments designed and implemented by student groups. All reactions were designed by high school students, with one round of intermediate feedback; the materials were prepared for nine student groups (30 students altogether) and reactions were implemented. Full details of each experiment, including students’ submitted reporters and all provided photos, are provided in supplemental materials. (B) Students constructed a cell-free lead sensor with a Mango III aptamer output and tested its activation against four input water sources, at two different times and two different temperatures. (C) Students constructed a cell-free copper sensor with a sfGFP output and tested it against three environmental water samples. (D) Students constructed cell-free fluoride sensors with three reporter outputs and tested the dose-response behavior for Mango, GFP, and XylE outputs. (E) Students constructed a cell-free lead sensor with RFP output and used it to measure lead in fruit juices.

We developed cascaded cell-free sensors for fluoride (using the *crcB* riboswitch [35]), copper (using the CueR aTF [69]), and lead (using the PbrR aTF [70]) (**Figure S4**). We also made negative and positive controls (in which o-T7 RNAP production is constitutively OFF or ON). When the lyophilized sensors were rehydrated with the respective orthogonal-promoter reporter plasmids (P_o-T7_ regulating expression of mRFP, sfGFP, Mango, or XylE), correct ligand-dependent induction was observed across all 80 possible combinations of five sensor plasmids, four reporter plasmids, and four inducers.

Armed with these designs, we asked nine groups of high school students who had already performed the LacI experiments to design and test their own biosensors. We specifically requested that they formulate hypotheses and devise positive and negative controls while allowing them to manipulate variables not tested in Modules 1-3. We provided each group the necessary reagents based on their experimental designs (e.g., reporter plasmids and inducers, freeze-dried sensor reactions containing the requested enriched aTFs/sensor plasmids) and simply requested photographs of the sensors at regular intervals after hydration to evaluate performance. The list of all the student-designed experiments is below in **Table 1**, and annotated pictures of sample experiments are in **Figure 4B**. The full student lab reports, including backgrounds and raw, uncropped photos are provided in the **Supplemental File 4**.

**Table 1.** A summary of student-led designs for Module 4 experiments.

| Group | Experiment |
| --- | --- |
| 1 | Detecting lead in vegetable and fruit samples |
| 2 | Measuring the limit of detection and leakiness of the lead sensor as a function of reporter output |
| 3 | Measuring the kinetics of the lead sensor with enzymatic reporter as a function of incubation temperature and lead concentration |
| 4 | Measuring the kinetics and signal of the fluoride sensor as a function of reporter output |
| 5 | Comparing the leak, dose response, and stability of response of the copper sensor with aptamer and enzymatic reporters |
| 6 | Measuring the dose-response of fluoride sensor with enzymatic reporter output |
| 7 | Detecting copper in environmental water samples |
| 8 | Measuring the specificity and kinetics of the lead sensor with the aptamer output at 26 and 37 °C |
| 9 | Measuring the limit of detection of the lead sensor with the enzymatic reporter |

Student success in Module 4 was varied, and many groups reported explicit sources of experimental error in their lab reports (*e*.*g*., contamination, loss of the lyophilized reaction pellet). However, this study provides a powerful proof-of-principle for the versatility of cascaded cell-free sensors, which allow students to detect any target molecule with any reporter output, while testing multiple induction conditions, temperatures, reporter plasmid concentrations, or sources of water samples. For instance, the reporter plasmid library could easily be extended to encompass fluorescent protein libraries [51] or enzymes that produce tactile or olfactory outputs [52]. These would be compatible with any sensed input, without needing to separately clone each inducible promoter.

### Large-scale implementation of biosensing education kits

To the best of our knowledge, all previous cell-free educational studies reported the results from hands-on experiments in professionally equipped laboratories. Our experiments in **Figures 2** and **3** were performed in high school classrooms, but these were still equipped with scientific instruments such as micropipettes. When many American high schools shut down in 2020 due to the COVID-19 pandemic and biology classrooms switched overnight to remote learning, we wondered if the intrinsic safety (i.e., no living cells) and thermal stability of freeze-dried cell-free sensors would allow them to be used for at-home experiential learning.

To test this possibility, we assembled the largest-scale distributed cell-free expression experiment to our knowledge. This consisted of 120 packaged Module 1 strip-tubes (720 total 20-µL CFE reactions, plus controls), which were individually packaged along with single-use, constant-volume pipettes, IPTG and water solution, and a desiccant card (**Figure 5A**). To test the long-term distributed stability of these reactions, we shipped the packages to two high schools in the Atlanta, Georgia metro area and presented the experiments to students along with accompanying curricula and pre- and post-lab surveys. At the time, students in both schools were in a hybrid setting, which meant that 32 students performed the experiments at home and 79 students completed them in a classroom. However, all students used plastic fixed-volume pipettes and incubated the reactions at room temperature. The success rate was measured qualitatively by photographs after 24 hours. These photographs, uncropped, are all available in **Supplemental File 5**.

**Figure 5.**
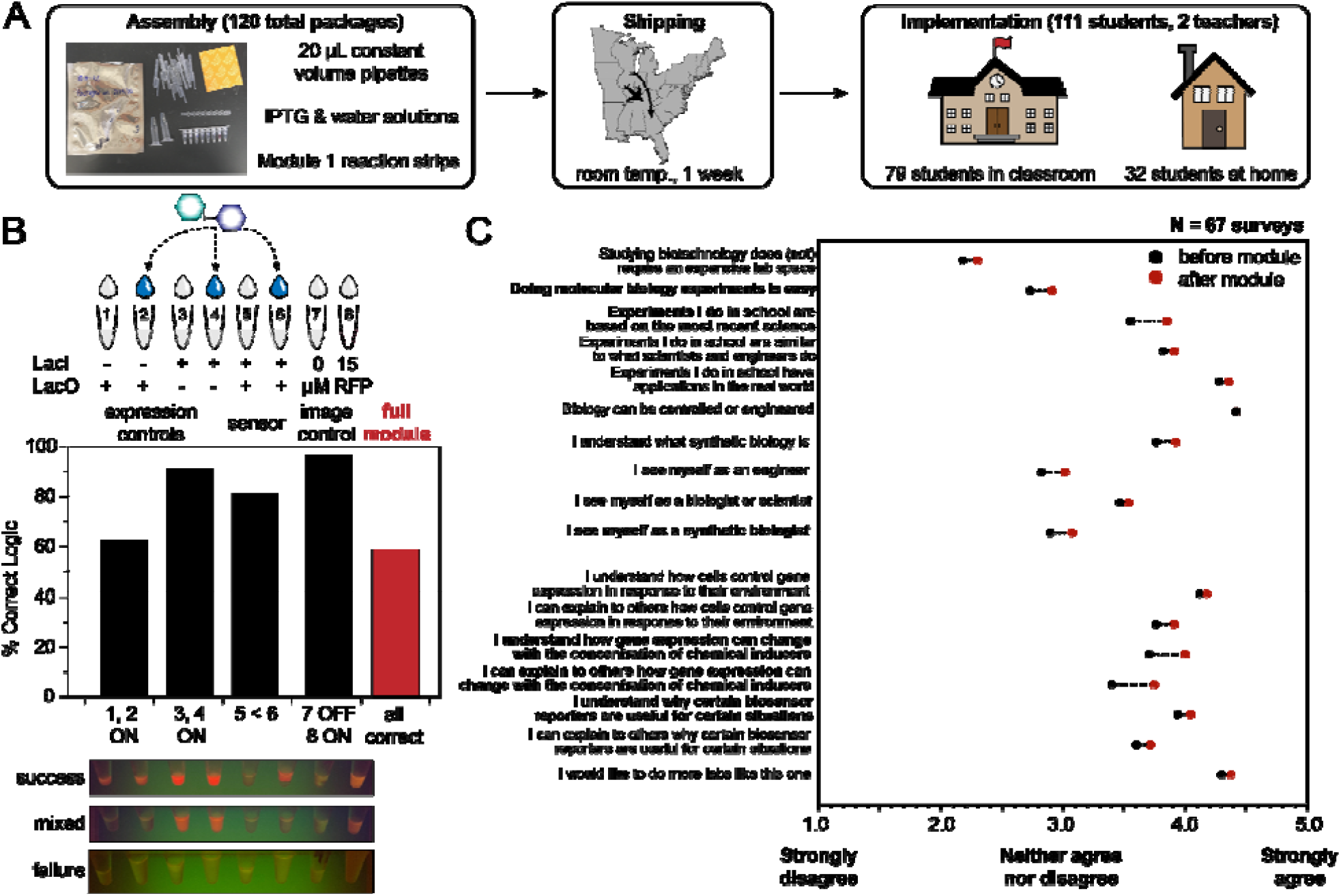
Large-scale implementation of remote learning biosensing module. (**A**) Overview of broad-scale deployment of Module 1, including outside-of-the-classroom learning. 120 Module 1 packages containing fixed-volume pipettes, stocks of IPTG and water, and the freeze-dried reaction eight-strips, were shipped from Northwestern University in desiccant packaging (photo shown) to two high schools in Georgia. 111 students participated, including 79 who performed the experiments in-person and 32 who ran the experiments at home. All students used the constant-volume pipettes to simulate a low-resource setting. (**B**) Overview of Module 1 success. Photos were taken after 24 hours of reaction incubation at room temperature in blue light (as available), and all tubes were scored as 0, 1, or 2 (“OFF”, “FAINT”, or “ON”) by two researchers. The average of each tube assignment was used to indicate a success rate for the whole module, with constitutive ON in tubes 1-4 and 8, OFF in tube 7, and a higher fluorescent signal in tube 6 than in tube 5. Around 60% of the reported photos showed complete success – that is, correct logic in every tube, though pipetting errors and lower-than-expected expression from the pLac-mRFP plasmid reduced the success rate relative to the small-scale local deployment in **Figure 2**. (**C**) Survey data overview, following the same analysis as in Figure 3, from the 67 remote students who completed pre- and post-lab survey for the module. Bolded statements indicate significant improvement (p < 0.05) in score.

The results (**Figure 5B**) from the distributed, large-scale Module 1 were less consistent than what we previously observed at the small scale (**Figure 2A**). Surprisingly low expression of mRFP was observed from both pairs of conditions that used the pLac-mRFP plasmid. In many cases (even in the absence of over-expressed LacI), tubes 1, 2, and 6 showed no visible activation. Among the subset of tubes where mRFP production was visible at endpoint, nearly all students did obtain constitutive expression from tubes 3 and 4 (J23119-mRFP) and observed IPTG-mediated induction in tube 6, compared to tube 5.

Overall, by scoring each tube’s brightness level as “OFF”, “FAINT”, or “ON”, we assessed around a 60% global success rate for the module, which we considered acceptably comparable to standard biology and chemistry classroom labs. We could not easily ascertain the origin for the failure mode at scale. However, when the remaining reactions were rehydrated by experienced biology teachers using micropipettes, the constitutive reactions activated well, and the fixed-volume pipettes also worked, although they were less accurate (**Figure S5A**). There were also several examples of student tubes that were clearly over-diluted with water or IPTG relative to the nominal pre-lyophilized volume. These effects do inhibit protein synthesis (**Figure 5B, Figure S5B**).

Considering the scale and purpose of the experiment, we did not repeat it. We instead emphasize that further work should be done to investigate the reproducibility of cell-free reactions at scale. This could include optimization of large-scale extract preparation, reaction assembly, and lyophilization prior to shipment as well as analyzing the impact of environmental fluctuations (e.g., temperature, humidity, *etc*.) between lyophilization and rehydration. Despite the mixed results from the experiment, students reported significant improvements in both “perspective” and “comprehension” fields on the survey after completing the experiment (**Figure 5C**). Importantly, students reported greater understanding of concentration-dependent behavior in biological systems as well as the benefits of different biological reporters.

## Conclusion

The cell-free biosensor modules developed here represent easy-to-use, low-cost, and distributable biology education labs that achieve learning outcomes through a 1-week curriculum with minimal equipment. After our research team designed and validated experiments based on the canonical *lac* operon, ∼20 high school students recapitulated laboratory data in their classrooms as they learned about the mechanisms, dose-responsive behavior, and common reporters for biosensors. Then, students undertook an engineering challenge to design, build, and test their own biosensor experiments using modular inputs (copper, fluoride, or lead ions) with modular outputs (fluorescent proteins, a fluorescent aptamer, or a colorimetric enzyme) for a more creative synthetic biology experience. Finally, we scaled up Module 1 (mechanism) for field testing with deployment to >100 students at schools 700 miles away and achieved ∼60% success across all modules using disposable pipettes in classrooms and home settings during the SARS-CoV-2 pandemic. The effectiveness of these modules was assessed through surveys before and after each set of experiments, revealing significant increases in several perspective and comprehension questions. Students reported increased understanding of biological sensors and reporters and felt that classroom labs were based on modern science after completing the modules.

Teaching principles of genetic regulation, biological sensing mechanisms, and field applications of synthetic biology in a hands-on fashion has the potential for significant impact. In total, we reached over 130 students across 3 high schools in 2 states with experiments implemented in classrooms or at home, not in well-equipped laboratories. We anticipate that further expansion of cell-free education modules like the ones described here will facilitate advances in hands-on STEM education by improving access to stimulate greater and earlier interest in biotechnology careers.

## Supplemental Files

### Supplemental Figures

Supplemental Figure 1. LacI-enriched extract titration.

Supplemental Figure 2. Reporter plasmid titrations.

Supplemental Figure 3. True/false survey questions.

Supplemental Figure 4. All cascade data pairs.

Supplemental Figure 5. Troubleshooting large-scale Module 1 distribution.

### Supplemental Files

Supplemental File 1. Uncropped raw photos and scores for Module 1 (Figure 2).

Supplemental File 2. Uncropped raw photos and scores for Module 2 (Figure 2).

Supplemental File 3. Uncropped raw photos and scores for Module 3 (Figure 2).

Supplemental File 4. Uncropped raw photos and scores for Module 1 (Figure 5).

Supplemental File 5. All lab reports for Module 4 (Figure 4).

Supplemental File 6. Lab curricula.

## Materials and Methods

### DNA assembly and purification

DNA was assembled using a mixture of commercial synthesis, PCR and blunt-end ligation, or isothermal (Gibson) assembly. pJBL7080 and pJBL7084 were synthesized by Twist Biosciences. pJBL7083 was synthesized by Gibson assembly using previously reported pJBL7010 and pJBL7072. pJBL7079 and pJBL7082 were synthesized by Gibson assembly using pJBL7080 and the constitutive expression cassettes as templates. pJBL7081 was assembled using overhang PCR and blunt-end ligation. The Module 4 plasmids were assembled using inverse PCR-blunt end ligation (all reporters: pJBL7056, pJBL7085, pJBL7086, pJBL7087) and Gibson assembly (all aTF expression cassettes and sensor plasmids, pJBL7093, pJBL7061, pJBL7062, pJBL7044, pJBL7045), with the exception of pJBL7063, which we could not successfully clone and was synthesized by Twist Biosciences. A list of strains, including descriptions and Addgene accession IDs, are presented below in **Table 2**. All annotated sequences are available on Addgene or by request.

**Table 2.** Summary of plasmids used in this manuscript. pLac refers to the *lacUV5* promoter. pT7 refers to the wild-type T7 promoter TAATACGACTCACATATA, and pAKSIRV refers to the orthogonal promoter variant TAATAC**CTGA**CACTATA, with the same 5’ UTR. AKSIRV refers to that T7 RNAP variant from [71]. J23119 is a consensus *E. coli* RNAP promoter [72].

| Plasmid ID | Plasmid Description | Addgene ID |
| --- | --- | --- |
| pJBL7044 | pT7-PbrR | 167215 |
| pJBL7045 | pT7-CueR | 167216 |
| pJBL7056 | pAKSIRV-sfGFP | 167225 |
| pJBL7061 | pPbr-AKSIRV T7 RNAP | 167230 |
| pJBL7062 | pCue-AKSIRV T7 RNAP | 167231 |
| pJBL7063 | J23119-fluoride riboswitch-AKSIRV T7 RNAP | 167232 |
| pJBL7079 | pLac-sfGFP | 167244 |
| pJBL7080 | pLac-mRFP1 | 167245 |
| pJBL7081 | pLac-Mango III | 167246 |
| pJBL7082 | pLac-XylE | 167247 |
| pJBL7083 | J23119-mRFP1 | N/A |
| pJBL7084 | pT7-LacI | 167248 |
| pJBL7085 | pAKSIRV-mRFP1 | 167249 |
| pJBL7086 | pAKSIRV-Mango III | 167250 |
| pJBL7087 | pAKSIRV-XylE | 167251 |

Plasmids were purified by Qiagen Midi Kits (ID 12143), quantified by NanoDrop, and stored long-term in water at −20 °C.

### Cell extract preparation

Cell extract was prepared to maximize expression from endogenous transcriptional machinery, as previously described [73] with a few modifications. For the laboratory-scale experiments in **Figures 1, 2**, and **4**, unenriched cell-free extract, 20 mL of a saturated overnight culture of BL21 Star (DE3) was inoculated into 1 L of 2X YT+P media (16 g/L tryptone, 10 g/L yeast extract, 5 g/L sodium chloride, 7 g/L potassium phosphate dibasic, and 3 g/L potassium phosphate monobasic adjusted to pH 7.2) and grown, shaking at 220 rpm at 37 °C, to optical density 3.0 (this required approximately 3 hours and 15 minutes for the base strain). The culture was decanted into 1 L bottles and cells were pelleted by centrifugation at 5,000 X g for 15 minutes at 4 °C. The cell pellets were washed once with 25 mL S30A buffer (14 mM Mg-glutamate, 60 mM K-glutamate, 50 mM Tris) and re-centrifuged for 10 minutes at 7,000 X g at 4 °C, following a report that one wash is sufficient to maintain good expression activity in the final extract [74]. The pellets were resuspended in S30A buffer at a ratio of 1 mL buffer/g pellet, transferred to 1.7 mL Eppendorf tubes, and sonicated on ice at 50% amplitude for 1 minute in six 10-second pulses using a QSonica Q125 small-tip probe. Immediately after sonication, 1 M dithiotreitol (DTT) was added to each tube to a final concentration of 3 mM. The lysate was clarified by centrifugation at 12,000 X g for 10 minutes at 4 °C. The top (supernatant) layer was removed, pipetted into a fresh tube, and incubated, shaking at 37 °C and 220 rpm, for 80 minutes for the ribosomal runoff reaction. After this time, the extract was centrifuged again at 12,000 X g for 10 minutes at 4 °C. 3 mL supernatant was transferred to a 10 kDa molecular-weight cutoff membrane and dialyzed against 600 mL S30B buffer (14 mM Mg-glutamate, 60 mM K-glutamate, 5 mM Tris, 1 mM DTT, pH 7.7) for 3 hours without exchange. After dialysis, the extract was transferred back to 1.7 mL Eppendorf tubes, clarified by one additional spin at 12,000 X g for 10 minutes at 4 °C, and the supernatant was removed, aliquoted, and flash frozen in liquid nitrogen.

For the enriched extracts (LacI, CueR, PbrR), a similar protocol was used. Chemically competent BL21 Star (DE3) cells were transformed with plasmids pJBL7084, pJBL7045, or pJBL7044 respectively, which encode the transcription factors under a T7 promoter. Overnight saturated cultures of these strains in LB were used to inoculate 1 L of 2X YT + P media and grown shaking, as before. Between optical density 0.4-0.5, 0.5 mM IPTG was added to induce protein synthesis. The strains were grown for 4-5 hours after induction and harvested as before. Extract was prepared identically, with the exception that no dialysis was performed on the enriched extracts. Instead, after the runoff reaction, the extracts were centrifuged, then directly aliquoted and flash frozen.

### Cell-free expression reaction

Cell-free gene expression was carried out as previously described^[73]^, in a mixture composed of 30 v/v% S12 extract; 8 mM magnesium glutamate, 10 mM ammonium glutamate, and 60 mM potassium glutamate; 1.2 mM ATP and 825 µM of CTP, GTP, and UTP; 34 mg/L folinic acid; 171 mg/L tRNA; 2.5 mM each amino acid; 30 mM phosphoenolpyruvate (PEP); 330 µM nicotinamide adenine dinucleotide (NAD); 270 µM coenzyme A; 4 mM potassium oxalate; 1 mM putrescine; 1.5 mM spermidine; 57 mM HEPES; plasmid DNA prepared with Qiagen Midi Kits; and the remainder water. Module 3 assays were performed with the addition of 1 mM catechol or 10 nM T01:biotin. Plasmid was added at 40 nM concentration in Modules 1-3 and was supplied in a 10X stock at 50 nM in Module 4.

### Lyophilization and storage

Prior to lyophilization, PCR strip tube flat caps (Axygen PCR-02-FCP-C) were punctured with a pin to create a hole, and PCR tube strips (Axygen PCR-02C) were placed into pre-chilled aluminum blocks on ice. Lyophilization was then performed by assembling the components of cell-free reactions as described above and placing them into pre-chilled PCR tube strips. Reaction tubes were then closed with the perforated PCR tube caps, submerged in liquid nitrogen, and transferred to a FreeZone 2.5 L Bench Top Freeze Dry System (Labconco). The reactions were then lyophilized for overnight with a condenser temperature of –85° C and 0.04 millibar pressure.

Unless rehydrated immediately, freeze-dried reactions were packaged as follows. About 2 - 5 strips of reactions (16 - 40 tubes total) were placed in a light-protective bag (Mylar open-ended food bags, Uline #S-11661) with a desiccant (Dri-Card Desiccants, Uline #S-19582). The reactions were then heat-sealed (Metronic 8-inch Impulse Bag Sealer, Amazon #8541949845) and stored in a cool, shaded area until usage.

### mRFP1 purification

mRFP1 purification was performed using a pET28c(+) expression plasmid pKJJ0062 (pT7-6XHis-TEV-mRFP). The sequence-verified plasmid was transformed into chemically competent Rosetta 2 (DE3) pLysS *E. coli*. A saturated overnight culture was inoculated into 1 L of LB media and grown at 37° C, then induced with 0.5 mM of IPTG at an optical density (600 nm) of ∼0.5 and grown for four additional hours at 37° C. Cultures were then pelleted by centrifugation at 5000 x g and were resuspended in lysis buffer (10 mM Tris-HCl pH 8, 500 mM NaCl, 1 mM TCEP, and protease inhibitor (cOmplete EDTA-free Protease Inhibitor Cocktail, Roche)). Resuspended cells were lysed on ice through ultrasonication, and insoluble material was removed by centrifugation. Clarified supernatant containing mRFP1 was then purified using His-tag affinity chromatography with a gravity column charged with Ni-NTA Agarose (Qiagen #30210). The elution from the gravity column was concentrated and buffer exchanged (25 mM Tris-HCl, 100 mM NaCl, 1mM TCEP, 50% glycerol v/v) using centrifugal filtration (Amicon Ultra-0.5, Millipore Sigma). Protein concentrations were determined using the Qubit Protein Assay Kit (Invitrogen #Q33212). The purity and size of the proteins were validated on an SDS-PAGE gel (Mini-PROTEAN TGX and Mini-TETRA cell, Bio-Rad). Purified proteins were stored at 4° C.

### Visual analysis of student reactions

Photographic student data was collected by imaging the reactions using a smartphone camera after approximately 24 hours incubation at room temperature. Photos were taken both in white light and in blue light using the miniPCR bio™ P51(tm) Molecular Fluorescence Viewer and were paired with (anonymized) tube identification numbers. We attempted but failed to quantitatively estimate RFP concentration relative to the 15 µM control tube from each photo, due to the poor image quality and unreliable lighting. Instead, we manually and qualitatively assigned each tube a value of “OFF”, “LOW”, and “ON”. Each tube in each photograph was ranked by at least two researchers to avoid bias. Examples of tubes ranked in each category are in **Figure 5**, and the full data set for each set of tubes is in the **Supplementary Files**. We then took the average of each qualitative measurement and used it to assign a success rate for the module.

### Survey analysis

Pre- and post-survey responses were paired for individuals according to their sample ID numbers, keeping student responses anonymous. Raw data sets were scrubbed to include only include paired, complete surveys, resulting in smaller sample sizes than the number of students who participated in the activities. For the binary questions, the asymptotic McNemar test with no continuity correction was used to assess statistical significance. For the categorical questions, the non-parametric Wilcoxon signed-rank paired test was used. This research was reviewed by the Northwestern Institutional Review Board Office and was determined to not be human research.

## Data Availability

Source data for all figures will be available in the Northwestern University Arch Institutional Repository or upon request.

## Acknowledgments

The authors acknowledge helpful conversations with members of the Jewett and Lucks labs. The authors gratefully acknowledge students at Evanston Township High School (Evanston, IL), Lambert High School (Suwanee, GA) and Centennial HS (Roswell, GA) for contributing data for this manuscript. This work is supported by the Army Research Laboratory and the Army Research Office (W911NF-22-2-0246), Army Contracting Command (W52P1J-21-9-3023), DTRA (HDTRA1-21-1-0038) the Department of Energy (DE-SC0023278), and the National Science Foundation (DGE – 2021900, CBET - 1936789). K.J.J. was supported in part by Northwestern University’s Graduate School Cluster in Biotechnology, System, and Synthetic Biology, which is affiliated with the Biotechnology Training Program (T32GM008449), by a Ryan Fellowship, and by a McCormick School of Engineering Terminal Year Fellowship. B.J.R. was supported by a National Defense Science and Engineering Graduate Fellowship (ND-CEN-017-095). G.A.R was supported by a National Science Graduate Research Fellowship (DGE-1842165). A.D.S. was supported in part by a Ryan Fellowship and a Presidential Fellowship. H.M.E. was supported in part by a National Defense Science and Engineering Graduate Fellowship (F-6669029987) and a Paul and Daisy Soros Fellowship. B.M.W. was supported by a National Science Graduate Research Fellowship (DGE-2234667).

