## Supplemental Information for "At-home, cell-free synthetic biology education modules for transcriptional regulation and environmental water quality monitoring"

a. Department of Chemical and Biological Engineering, Northwestern University, Evanston, IL 60208, USA
b. Chemistry of Life Processes Institute, Northwestern University, Evanston, IL 60208, USA
c. Center for Synthetic Biology, Northwestern University, Evanston, IL 60208, USA
d. Center for Water Research, Northwestern University, Evanston, IL 60208, USA
e. Interdisciplinary Biological Sciences Graduate Program, Northwestern University, Evanston, IL, 60208, USA
f. Lambert High School, Suwanee, GA 30024, USA
g. Centennial High School, Roswell, GA 30076, USA
h. Fulton County Schools Innovation Academy, Alpharetta, GA 30009, USA
i. Evanston Township High School, Evanston, IL 60201, USA
j. Robert H. Lurie Comprehensive Cancer Center, Northwestern University, Chicago, IL 60611, USA
k. Simpson Querrey Institute, Northwestern University, Chicago, IL 60611, USA
l. Department of Bioengineering, Stanford University, Stanford, CA 94305

^#^ These authors contributed equally to this work.

* Corresponding authors

**Correspondence**

Julius B. Lucks,

Michael C. Jewett,

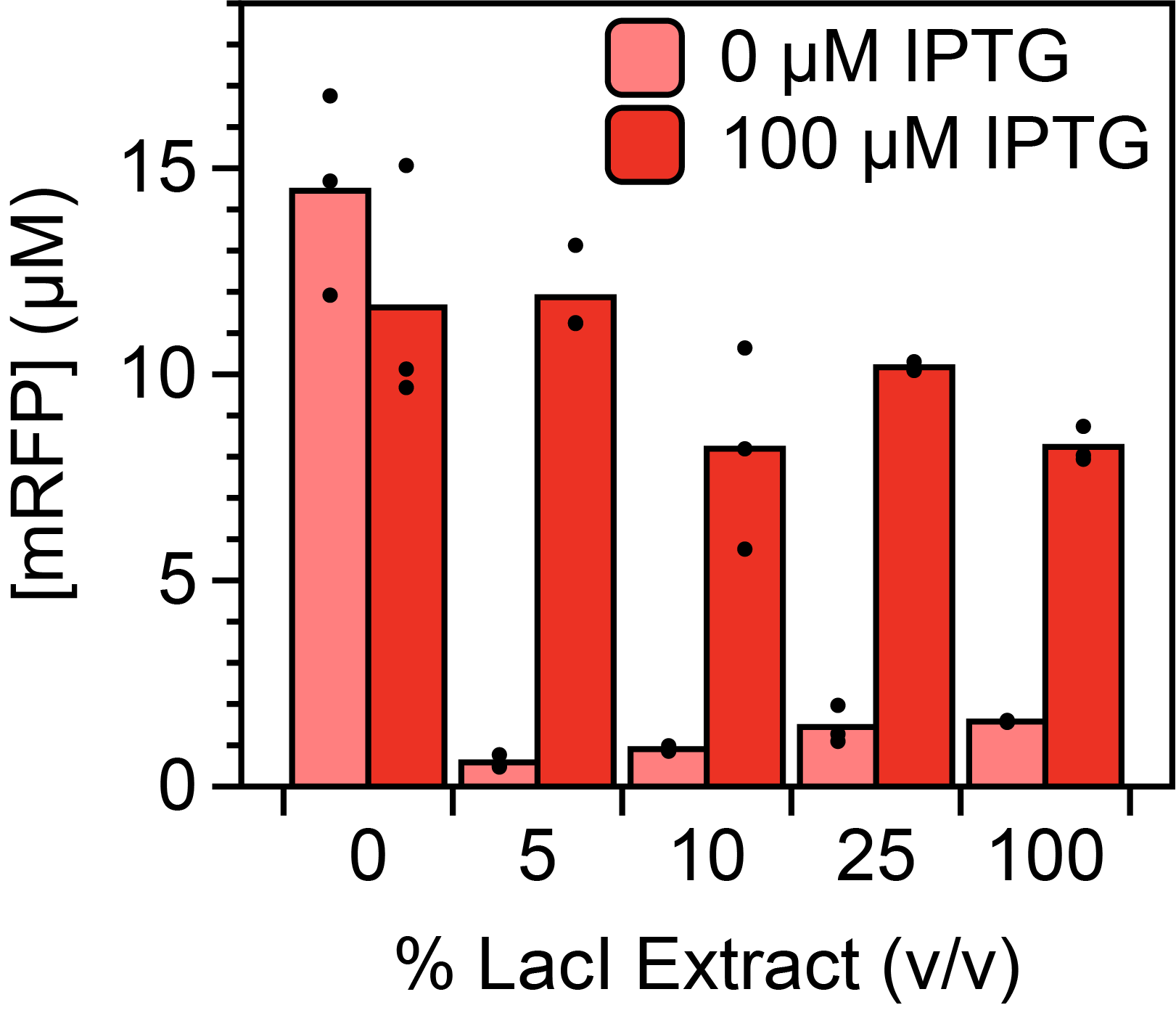

**Figure S1. LacI-enriched extract titration.** CFE reactions with 20 nM pLac-mRFP1 plasmid were constructed with varying proportions of unenriched extract and LacI-enriched extract. 5% LacI extract was selected for Modules 1-3 to have maximum signal and minimal leak.

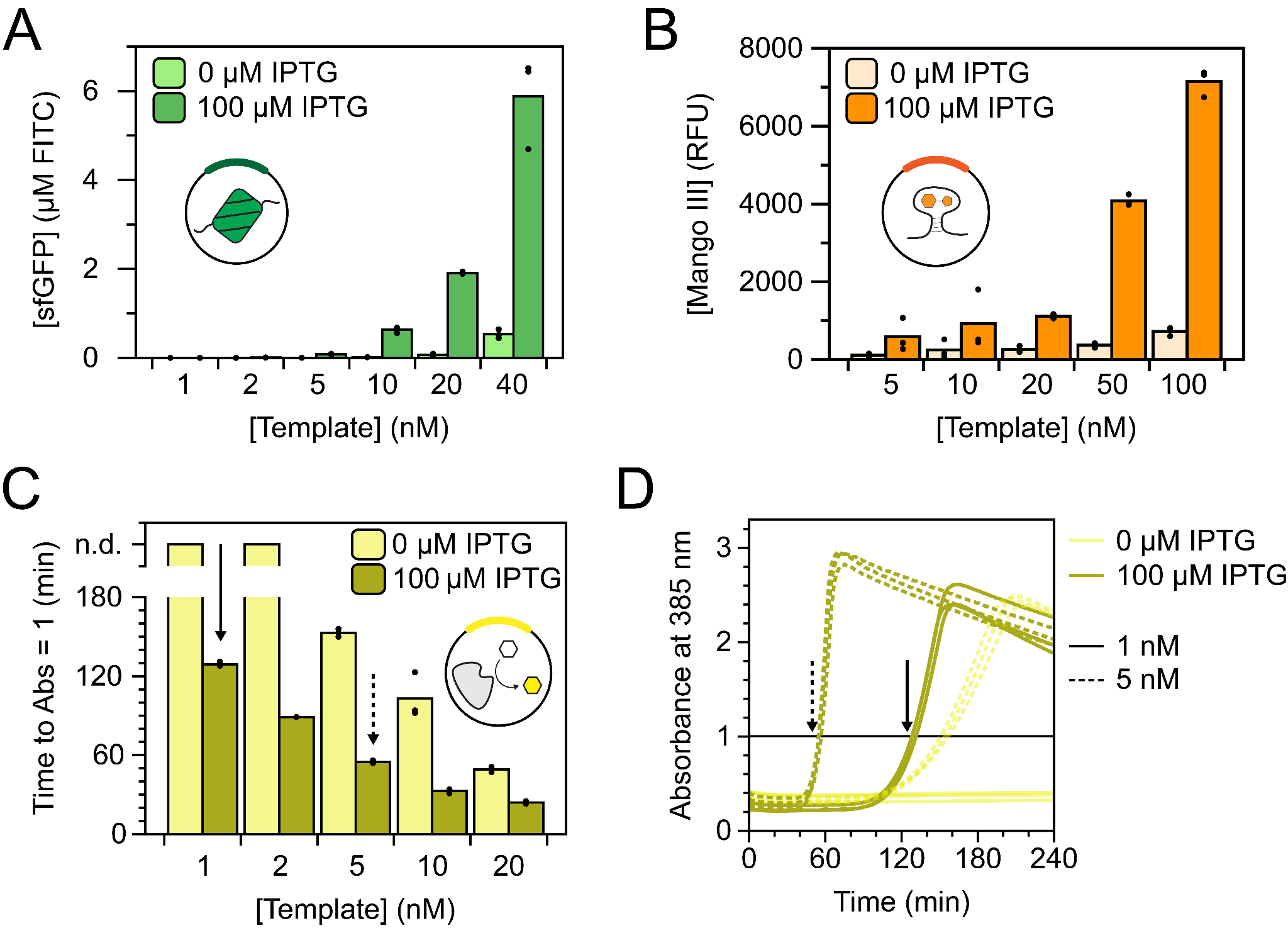

**Figure S2. Reporter plasmid titrations.** CFE reactions with 5% LacI-enriched extract were constructed with varying concentrations of expression templates for sfGFP (A), Mango III aptamer (B), and XylE (C) to maximize signal and minimize leak. The data in panel C represent time to a threshold absorbance of 1.0, linearly interpolated from absorbance measurements every two minutes over four hours, where the lower value indicates a faster time-to-result. The corresponding kinetic traces for two of the concentrations tested (1 nM and 5 nM) are shown in panel D, and the threshold absorbance of 1.0 is indicated with a solid black line.

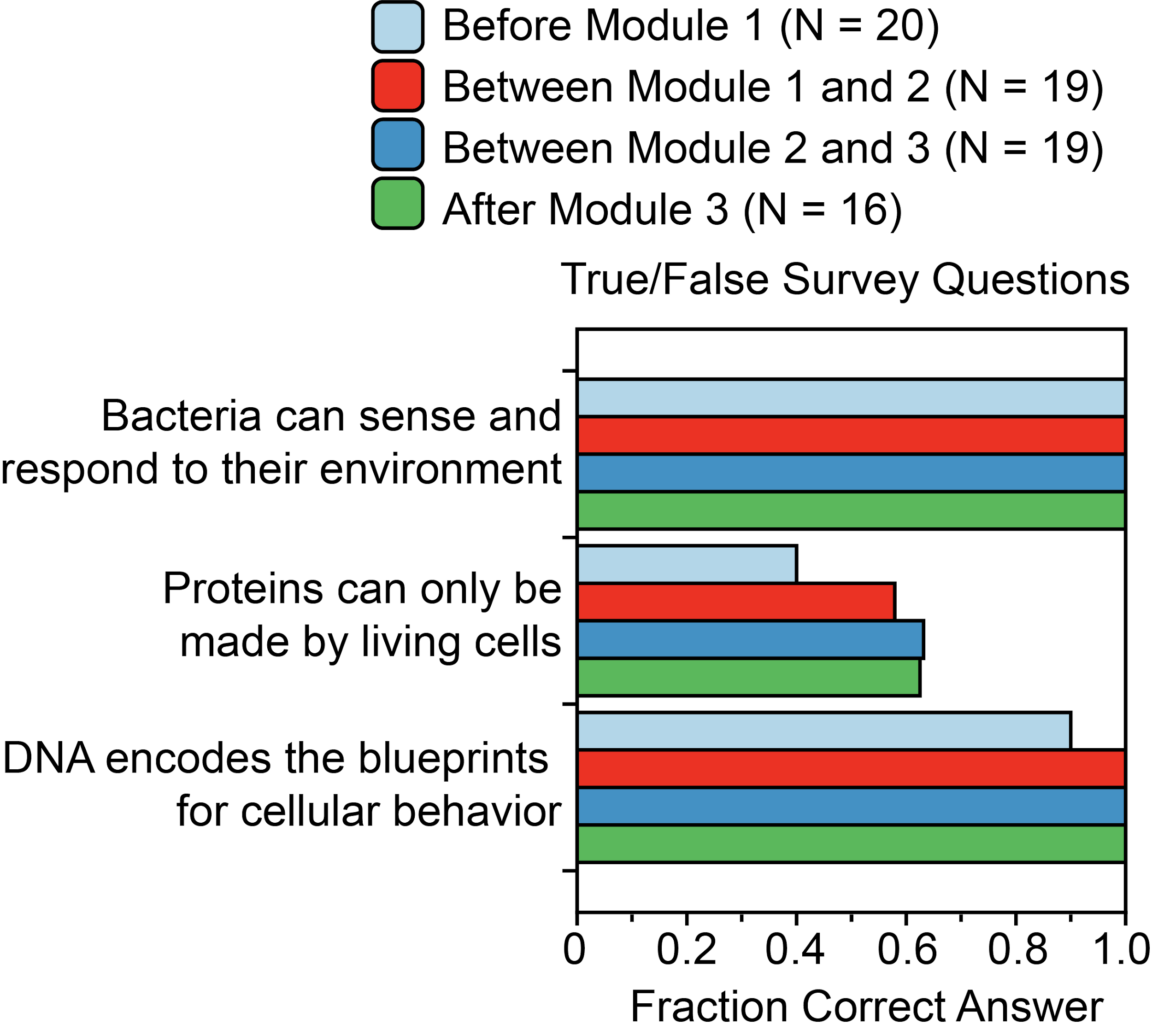

**Supplemental Figure S3. True/False survey questions.** Students at Evanston Township High School completed true/false questions along with the perspective and comprehension questions in the survey. Most students correctly answered the first and third questions prior to completing the experiment modules, but the curricula did result in more students correctly answering the second question.

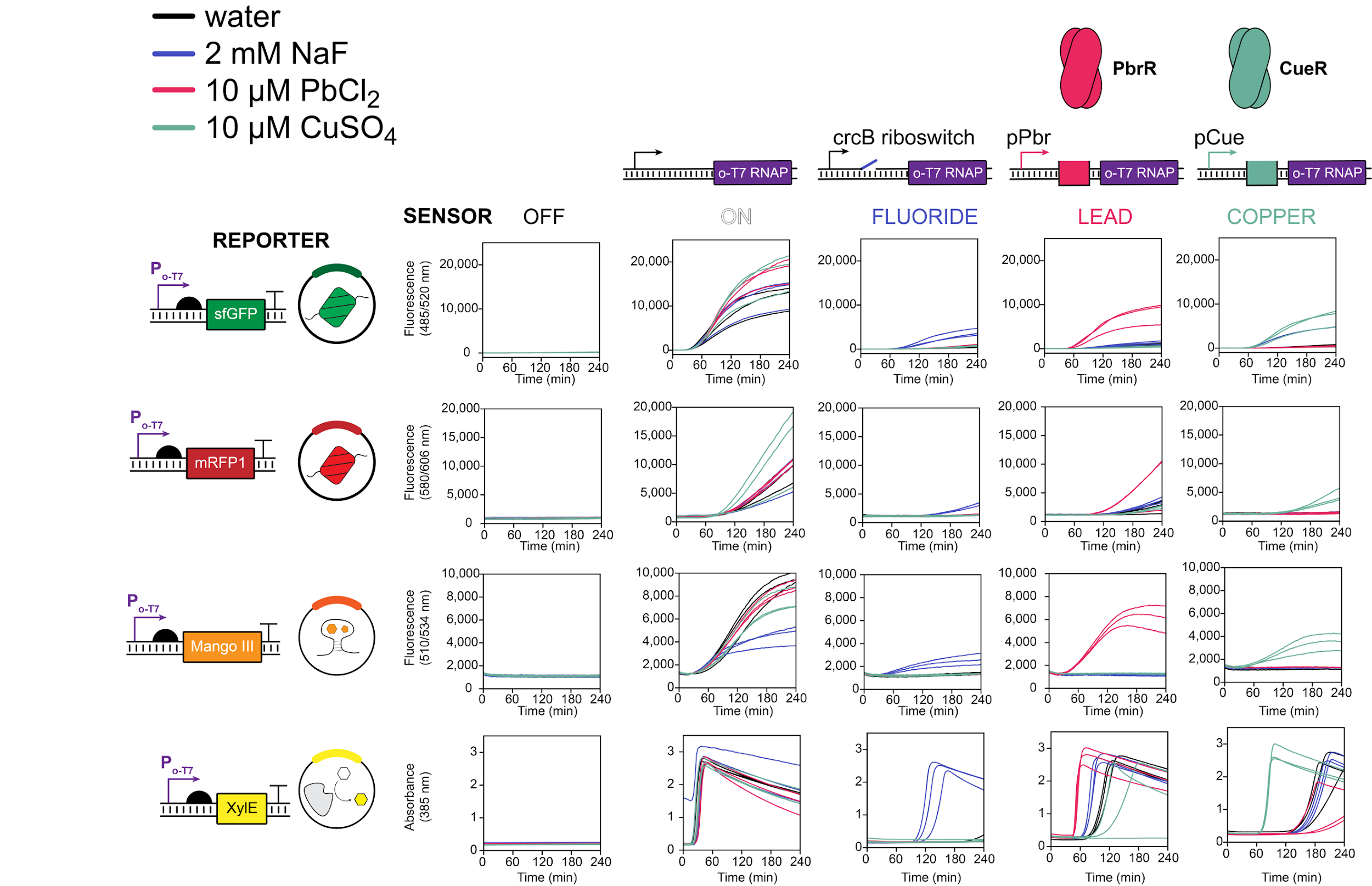

**Figure S4. All cascade data pairs.** Transcriptional cascades were run for all combinations of sensor and reporter plasmids available to students for Module 4 to verify modularity and effective construction of many unique sensing reactions. Plotted data represent the individual trajectories from three replicates of every possible lyophilized sensor reaction (5 sensor conditions rehydrated with each of 4 reporter plasmids and each of 4 inducer conditions).

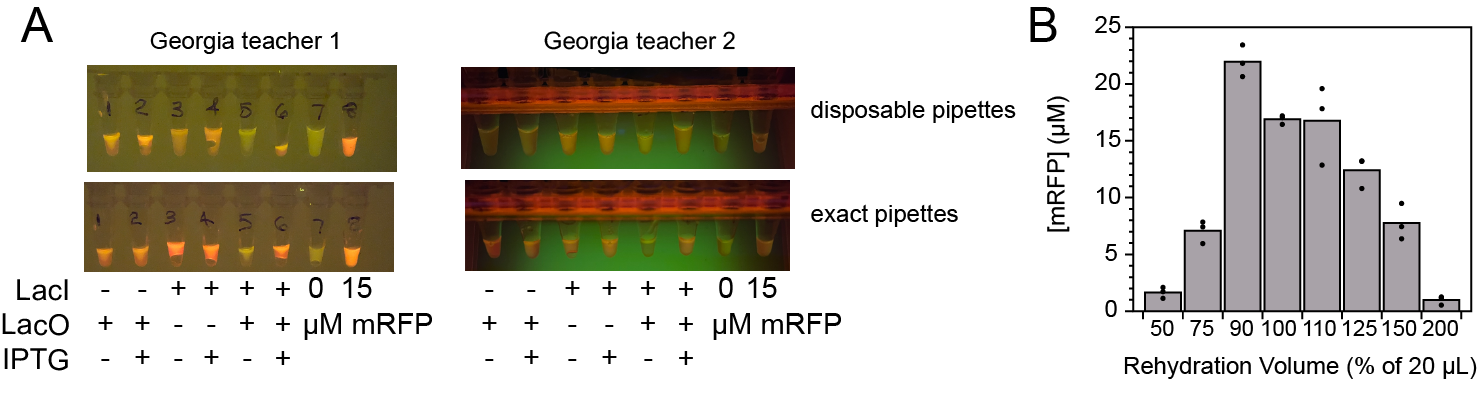

**Figure S5. Troubleshooting large-scale Module 1 distribution.** (A) Before the Georgia students performed Module 1, their teachers rehydrated the remaining reactions and observed results consistent with laboratory data (2 strip-tube replicates from each school). (B) The rehydration volume is a key consideration for CFE output. Plotted data represent the endpoint mRFP yield after four-hours when a 20 µL lyophilized IPTG-sensing reaction is rehydrated with varying volumes of a 100 µM IPTG stock. Significantly over- or under-diluting the CFE reactions can result in repressed protein synthesis. This is one of the main drawbacks of disposable pipettes that are less accurate and reliable than laboratory micropipettes.
